# The PhoPQ two-component system is the major regulator of cell surface properties, stress responses and plant-derived substrate utilisation during development of *Pectobacterium versatile*-host plant pathosystem

**DOI:** 10.1101/2020.04.24.060806

**Authors:** Uljana Kravchenko, Natalia Gogoleva, Anastasia Kolubako, Alla Kruk, Julia Diubo, Yuri Gogolev, Yevgeny Nikolaichik

**Affiliations:** Department of Molecular Biology, Belarusian State University, Minsk, Belarus; Kazan Institute of Biochemistry and Biophysics, Federal Research Center “Kazan Scientific Center of RAS”, Kazan, Russia

## Abstract

The PhoPQ two-component system, originally described in pectobacteria as PehRS, was previously shown to regulate a single gene, *pehA*. Using an insertional *phoP* mutant of *Pectobacterium versatile*, we demonstrate that PhoP controls a regulon of at least 116 genes with a large fraction of regulon members specific for pectobacteria. The functions performed by the PhoP controlled genes include transport and metabolism of plant-derived carbon sources (polygalacturonate, arabinose and citrate), modification of bacterial cell envelope and stress resistance. High concentrations of calcium and magnesium ions were found to abolish the PhoPQ-dependent transcription activation. Reduced PhoP expression and minimisation of PhoP dependence of regulon members’ expression in the cells isolated from rotten potato tuber tissues suggest that PhoPQ system may adjust expression levels of multiple virulence-related genes during the course of *P. versatile*-host plant pathosystem development.

## Introduction

*Pectobacterium* spp. are pectinolytic bacteria that cause soft rot and other diseases in a variety of plants. Until recently, members of this genus were often considered to be broad host range pathogens. However, genomic studies revealed significant diversity within this group, resulting in the ongoing subdivision of the previously described species and elevation of subspecies to species level. This resulted in better separation of the related strains more or less according to their environmental preferences. *Candidatus Pectobacterium maceratum* was suggested as a new name for a group of strains isolated from potato and cabbage (1). Recently, more isolates from various sources, including *Chrysanthemum, Iris* and water, were united with *Ca. P. maceratum* strains and renamed as *Pectobacterium versatile* (2). The *P. versatile* taxon includes two strains which have been well characterised experimentally, Ecc71 and SCC1. The 3-2 strain used in this work also belongs to this species.

*P. versatile* (*Pve*) strains can infect various plant hosts, and the same strain can cause diverse symptoms while infecting distinct plant tissues (e.g. stem blackleg and tuber soft rot in potato). Pectobacteria are also known to proliferate stealthily for a while in the vascular tissues before switching to a brute force attack of surrounding tissues via the massive secretion of plant cell wall hydrolases (3). Such versatility of *Pectobacterium* spp. virulence strategies must be carefully controlled at the transcriptional level. Yet, only a small fraction of about 300 transcription factors have been characterised in this genus. These include pectinolysis and exoenzyme regulators KdgR (4), RexZ (5) and GacA (6,7), the alternative sigma factor HrpL – the activator of the type III secretion system and its substrate genes (8), motility regulators HexA (9) and FlhDC (10), pectin lyase regulators RdgA and RdgB (11) and quorum sensing regulators ExpR and VirR (12–15).

PehR was originally described as a transcriptional activator of the *pehA* gene encoding endopolygalacturonase, a major virulence factor in soft rot bacteria (16). PehR is a response regulator that forms a two-component sensory system (TCS) with the membrane histidine kinase PehS. In the SCC3193 strain, which is currently classified as *Pectobacterium parmentieri*, inactivation of either *pehR* or *pehS* resulted in reduced virulence and Ca^2+^ was reported as the ligand detected by the PehS sensor (16,17). It was also suggested that the PehRS system is responsible for the decrease of the polygalacturonase and increase of the pectate lyase activities in response to Ca^2+^ released from the degraded cell walls (17).

The pectobacterial PehRS two-component system received very little attention since its discovery two decades ago. However, much information has been accumulated about the orthologous system called PhoPQ in other bacteria from the order *Enterobacetrales*. In *Dickeya dadantii*, which belongs to the *Pectobacteriaceae* family together with *Pve*, PhoPQ TCS was shown to control pellicle formation, motility, resistance to cationic antimicrobial peptides and expression of pectate lyases in response to Mg^2+^ concentration (18,19). The PhoPQ system has been thoroughly characterised in *Salmonella* and to a lesser extent in few more *Enterobacteriaceae* species including *Escherichia coli* and *Yersinia* spp. (see (20) for a review). The PhoQ sensor histidine kinase activates PhoP by phosphorylation in response to several stimuli, including low Mg^2+^ concentration (21), cationic peptides (22), acidity (23) and high osmolarity (24). PhoP binds to direct repeats with a consensus gGTTTA which seems to be well conserved in enterobacteria (25,26). The regulon composition nevertheless varies widely even between closely related bacteria. In *Salmonella enterica*, over a hundred genes are regulated by PhoP, but only three of them (*phoP, phoQ* and *slyB*) are always under PhoP control in other enterobacteria. Some regulon members are shared by several species, but most PhoP controlled genes are species and even strain-specific (25). In summary, PhoP is a global regulator controlling diverse, but always large, regulons in *Enterobacterales*.

Despite the obvious importance of PhoP in *Enterobacterales*, its regulon was not studied so far in *Pectobacterium spp*. To date, *pehA* remains the only known PhoP (PehR) target in these bacteria. The global mode of regulation reported for PhoP in other species strongly suggests that in pectobacteria it is likely to control many genes in addition to *pehA*. Moreover, in the recent work on *P. atrosepticum* transcriptome profiling, we have noticed downregulation of *phoPQ* (27). Expression level changes *in planta* combined with a potentially large regulon suggested that PhoP might be an important regulator of virulence properties in pectobacteria.

In this study we demonstrate that PhoP regulon of *Pve* includes at least 116 genes and incorporates a significant number of *Pectobacterium*-specific genes. We also try to distinguish the regulon parts directly and indirectly controlled by PhoP and discuss the implications of regulon composition for the regulation of *Pectobacterium* virulence.

## Experimental procedures

### Bacterial strains and growth conditions

*Pve* strain JN42 (28) is a spontaneous rifampicin-resistant derivative of the wild type isolate 3-2, originally described as *Erwinia carotovora*. The *Escherichia coli* strain XL-1 Blue (29) was primarily used for plasmid construction and *E. coli* strain BW 19851 (30) was used for conjugational transfer of suicide vector pJP5603 (31) derivatives into *Pve. Pve and E. coli* were routinely grown in lysogeny broth (LB) or minimal medium at 28 °C and 37 °C respectively.

Two minimal media were used throughout this work. Minimal medium A (MMA) was composed of K2HPO4 (10,5 g/l), KH2PO4 (4,5 g/l), (NH4)2SO4 (1 g/l), sodium citrate (0,43 g/l) and 0.5% glycerol. MgSO4 was added to MMA to the final concentration of 0.5 mM. Sodium polypectate (Sigma) or L-arabinose were added to the final concentrations of 0.5% and 0.2% when necessary. To avoid precipitation, the effects of different divalent cation concentrations were studied in minimal medium N (MMN) (32) containing KCl (5 mM), (NH4)2S04 (7.5 mM), K2S04 (0.5 mM), KH2PO4 (1 mM), and Tris-HCl (0.1 M), pH 7.4 and 0.5% glycerol.

Antibiotics were used at the following concentrations (µg/ml): ampicillin, 100; gentamycin, 10; rifampicin, 25.

### Construction of the *phoP* mutant and complementation plasmid

To construct a *phoP* mutant of *Pve*, the *phoP* gene sequence was PCR amplified from *Pve* 3-2 with phoPf and phoPr primers (Table 1) and its internal NdeI-PvuII fragment (225 bp) was cloned into the suicide vector pJP5603. The resulting plasmid was mobilized into *Pve* from *E. coli* BW 19851 and *Pve* crossover clones were selected on kanamycin (20 μg/ml) containing plates. *Pve* disruption was confirmed by PCR with combinations of primers to *phoP* and suicide vector sequences (phoPf-phoPr, phoPf-pjp2 and phoPr-pjp1, Table 1).

**Table 1.**
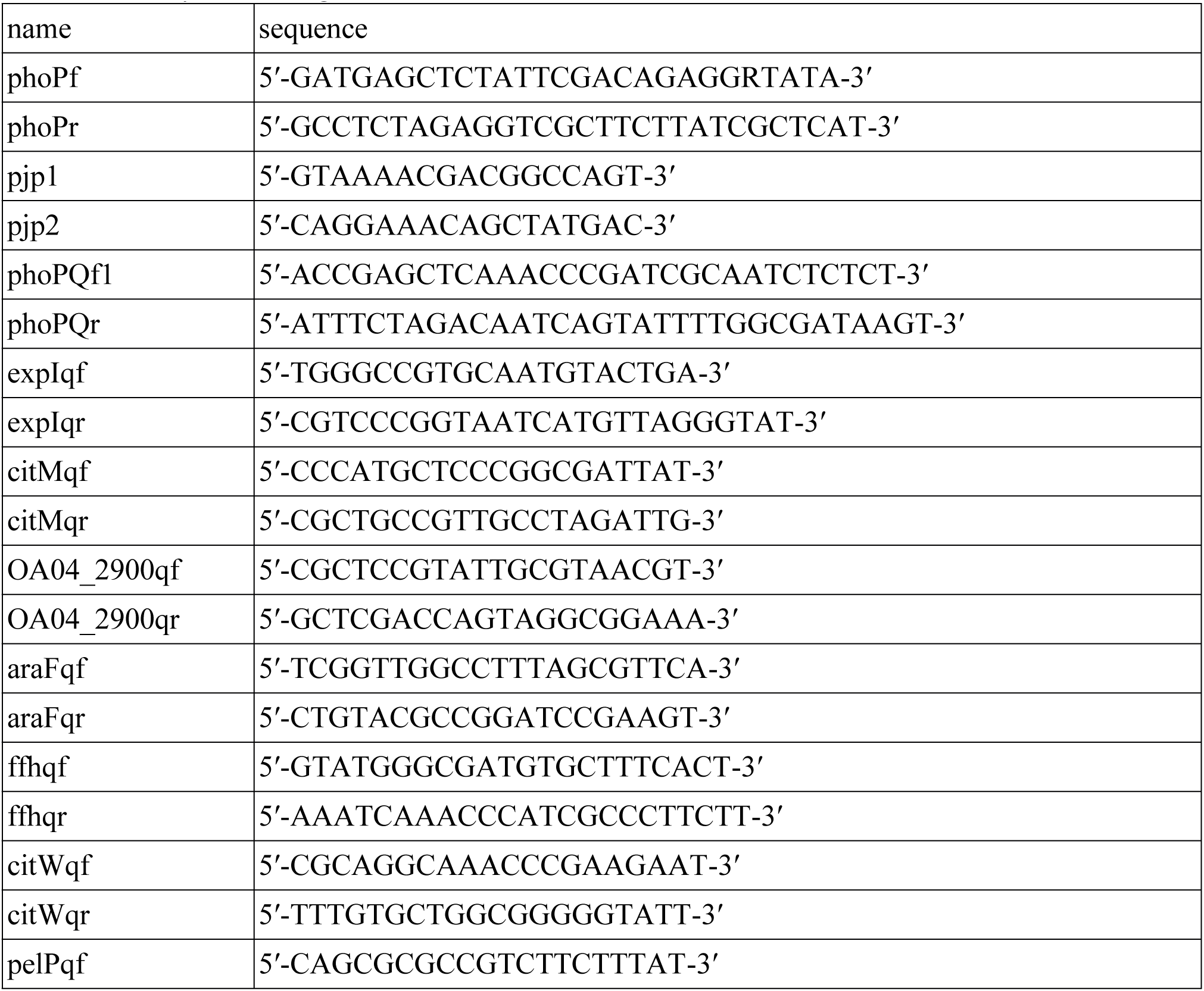
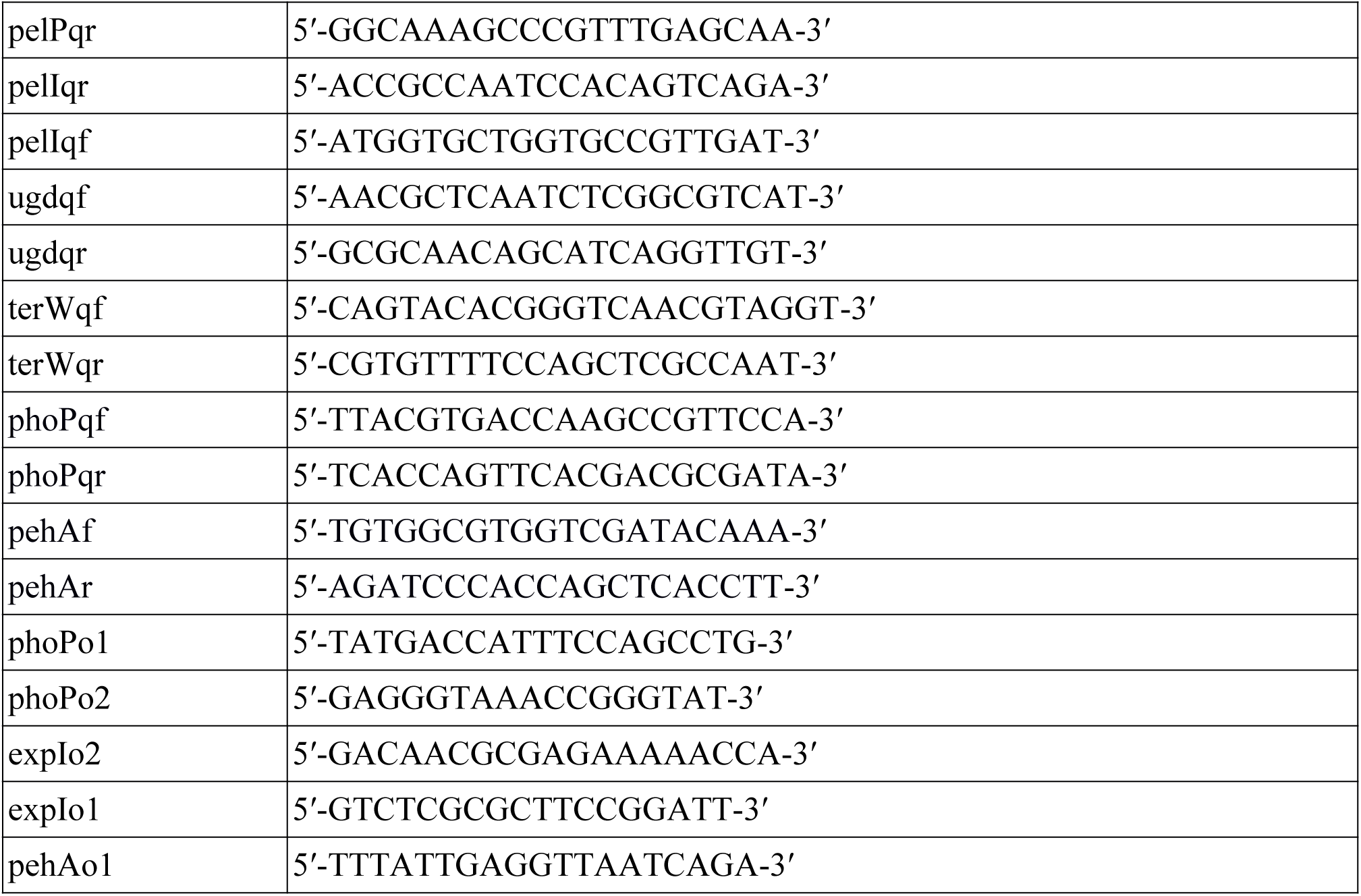
Synthetic oligonucleotides.

The plasmid expressing *phoP* and *phoQ* was constructed by cloning the PCR fragment amplified with the Tersus DNA polymerase (Evrogen) and the phoPQf1 and phoPQr primers into the low copy number vector pZH449.

Characteristics of the plasmids used in this work are specified in Table 2.

**Table 2.** Plasmids.

| Plasmid name | Characteristics | Source/reference |
| --- | --- | --- |
| pJP5603 |  | (31) |
| pJP5603::'phoP' |  |  |
| pJQ200KS | Gm<sup>r</sup> *lacZα* *mob* oriV<sub>p15A</sub> | (33) |
| pZH449 | 1.8 kb ClaI-HindIII deletion of pJQ200KS which removed *sacB* | This work |
| pPQ | pZH449::phoPQ | This work |
| pET42a |  | Novagen |
| pET42a::phoP |  | This work |

### RNA extraction, cDNA library preparation and sequencing

To extract total RNA, *Pve* cells from the aliquots of cell suspensions or macerated potato tissue were fixed by adding phenol/ethanol (1/20 v/v) solution to 20% and keeping on ice for 30 min. The fixed cells were harvested (8000 g, 5 min, 4 °C) and resuspended in 1 mL of ExtractRNA Reagent (Evrogen, Russia) and the subsequent procedures were performed according to manufacturer’s instructions. Residual DNA was eliminated by treatment of RNA samples with DNAse I (Thermo Fischer).

For RNA-Seq, total RNA was processed using Ribo-Zero rRNA Removal Kit (Gram-Negative Bacteria) (Illumina) and NEBNext Ultra Directional RNA Library Prep Kit for Illumina (NEB), according to manufacturer’s instructions. The quality and quantity of the cDNA libraries during processing before sequencing were monitored using the Agilent 2100 Bioanalyser (Agilent) and CFX96 Touch Real-Time PCR Detection System (Bio-Rad). Sequencing was conducted by a HiSeq 2500 Sequencing System (Illumina, USA) at Joint KFU-Riken Laboratory, Kazan Federal University (Kazan, Russia). Each type of the libraries (wild type and *phoP* mutant) was sequenced in four biological replicates.

### RNA-Seq data analysis

The coding sequences of *Pve* 3-2 genome were used as a reference (GenBank accession CP024842). The obtained read sequences corresponding to the Pve genome can be accessed from NCBI’s BioProject under the accession number PRJNA627079. Read pseudo-alignment and transcript quantification was performed using the alignment-free kallisto tool (34). The analysis of the differentially expressed genes (DEGs) was carried out with the edgeR package (35). Genes with fold-change > 2 and significant differences in expression levels (FDR < 0.05) were considered as DEGs.

### Gene expression analysis by qPCR

The total RNA from *Pve* cells was extracted, treated with DNAse and quantified as described above. 1 μg of RNA was used for cDNA synthesis using RevertAid reverse transcriptase (Thermo Scientific) according to the manufacturer’s instructions. Two μl of 5-fold-diluted cDNA were used as the template for qPCR.

qPCR was performed using a SYBR Green I - containing master mix. Primers for target and reference genes (Table 1) were designed using primer3 (36) and checked for specificity with Primer-Blast (37). PCR was performed under the following conditions: 95 °C for 2 min, followed by 45 cycles at 94 °C for 10 s and 60 °C for 60 s. After that, melting curve analysis was performed in the temperature range of 60–90 °C. The reactions were run and changes in fluorescence emission were detected using a DT-96 quantitative PCR system (DNA Technology, Russia). The amount of fluorescence was plotted as a function of the PCR cycle and converted to Cp values using RealTime_PCR Software (DNA Technology, Russia). The amplification efficiency for all primers was determined using a dilution series of genomic DNA. Additional controls included the omission of reverse transcriptase to measure the extent of residual genomic DNA contamination and template omission. The *ffh* and *gyrA* genes, the transcript levels of which were confirmed by the geNorm software (38) to be stable under the applied experimental conditions (data not shown), were used for normalization of the expression of the target genes. At least four biological replicates were performed for each measurement. Relative expression levels and error estimates were calculated using the REST 2009 v. 2.0.13 software (Quiagen). In all cases, the mean values are shown with 95% confidence interval.

### Transcription factor binding site inference and correction of genome annotation

ChIPmunk (39) was used for the inference of PhoP binding sites within the regulatory regions of the differentially expressed genes. The ChIPhorde version of the algorithm was run, followed by manual filtering of putative operators according to their scores and positions relative to transcription initiation sites. TFBS positions in the context of RNA-seq coverage were visualised by SigmoID (40). Wig files with RNA-seq coverage data were generated by Rockhopper (41).

Inference of operator motifs for other TFs was done with the modified algorithm of Sahota and Stormo (42) as implemented in SigmoID version 2.0 (https://github.com/nikolaichik/SigmoID). SigmoID was also used for scanning the *Pve* genome for operator sequences matching known or new motifs. Positions of critical residues required by this algorithm were determined by the interaction service of NPIDB (43). Alignment of the DNA binding domains was done with hmmalign from the HMMER3 package (44).

Correction of annotation for PhoP controlled genes was done using search and editing capabilities of SigmoID (40). The updated annotation of the *Pve* 3-2 genome was submitted to GenBank under the same accession number.

### Virulence assays

Bacteria were grown overnight on solid LB plates, washed off with 0.85% NaCl solution, centrifuged briefly and resuspended in the same solution, after which cell suspension densities were adjusted to achieve the cell doses 5·10^5^ (high inoculum dose) or 1·10^5^ (low inoculum dose) per inoculation site. Potato (*Solanum tuberosum*) cv Zhuravinka tubers were inoculated with 10 µl of cell suspensions using a 200 µl pipette tip close to stolon end of the tuber. The inoculation sites were wrapped in parafilm and tubers were kept in plastic bags for 48 h at 28 °C.

For Chinese cabbage inoculation, cell suspensions were prepared the same way, but their cell densities were adjusted to the values 1·10^5^ (high inoculum dose) or 1·10^3^ (low inoculum dose) per inoculation site. Chinese cabbage leaves were soaked in the sodium hypochlorite solution (70-85 g/ l of active chlorine) and then flushed with sterile deionised water. Rectangular sections were cut out from the base of each leaf. The obtained sections were soaked in the 96% ethanol and then flamed. The sections were inoculated with 10 µl of cell suspensions using a 200 µl pipette tip. The samples were placed in the sterile Petri dishes and incubated for 48 h at 28 °C.

At least ten tubers or leaf sections were used per each experimental condition.

## Results and discussion

### Inactivation of *phoP* alters *P. versatile* virulence

To examine the role of PhoP in *P. versatile* interaction with host plants, we compared the properties of the *phoP* mutant strain with its wild type parent and a complementation strain. Since *phoP* is the first gene of the *phoPQ* operon and a polar effect of *phoP* inactivation on the expression of *phoQ* was expected, complementation was achieved by a plasmid carrying the whole *phoPQ* operon together with the full regulatory region.

In Chinese cabbage virulence tests carried out at low inoculum doses (10^3^ cells), the mutant strain caused less damage than the wild type strain, and observed virulence defect was fully complemented by the plasmid expressing *phoPQ* (Figure 1a,b). We could not detect reproducible differences between the strains at higher inoculum doses (data not shown). In potato tubers, we could reliably induce infection starting with much higher inoculum dose (10^5^ cells) and no difference between the strains was observed. However, the *phoP* mutant generated 50% more macerated tuber tissue with a dose of 10^5^ cells (Figure 1c). In this case, the *phoP* mutant caused more tissue maceration than the wild type strain. Since low and high inoculum doses somewhat mimic the early and late stages of infection development, virulence tests results indicate possible importance of the PhoPQ two-component system through the whole course of the infection process.

**Figure 1.**
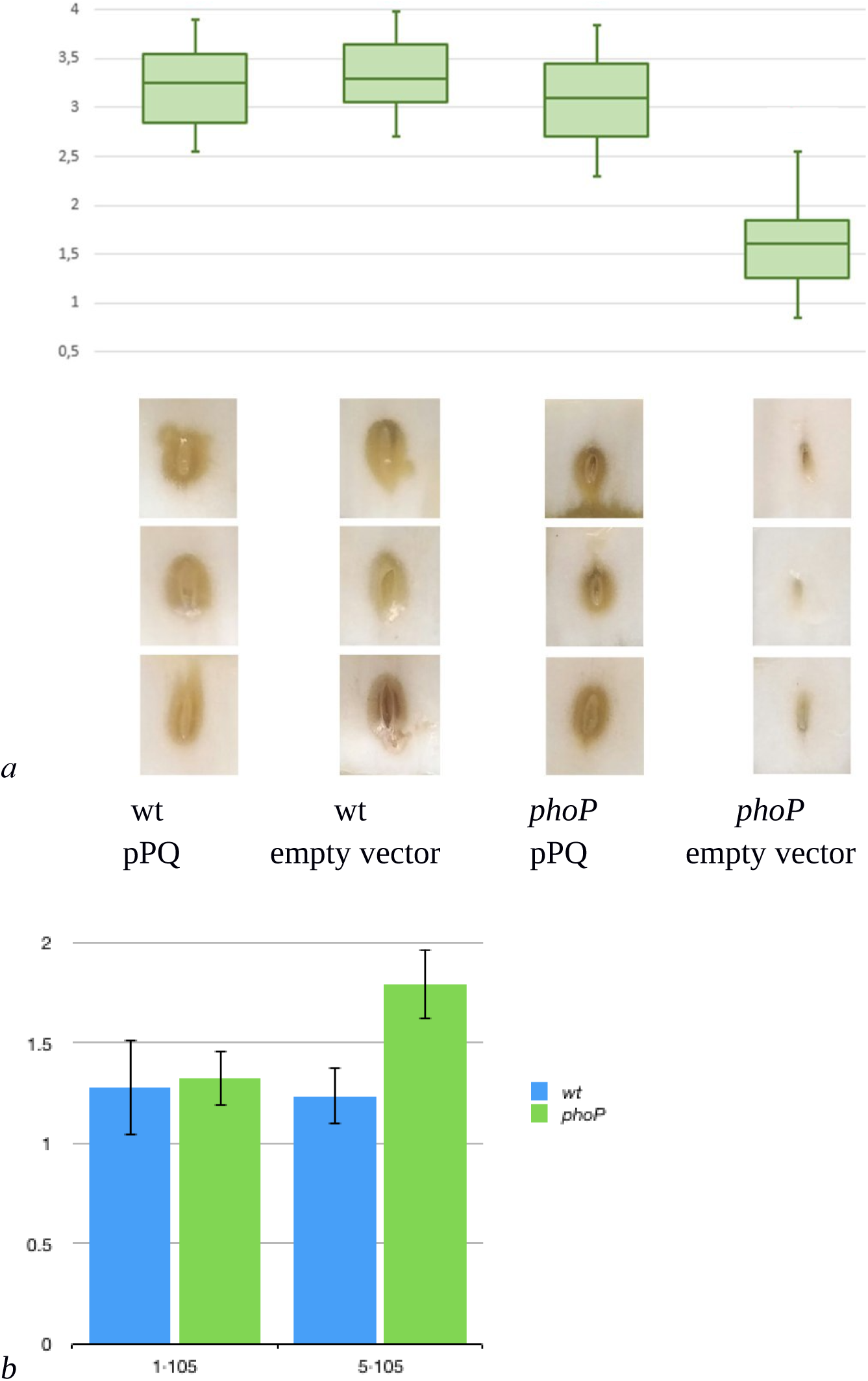
Effect of *phoP* inactivation on *Pve* virulence. a) Boxplot representation of disease symptoms induced in Chinese cabbage leaves by *Pve* strains JN42 (wild type) and UK1 (*phoP* mutant) carrying either an empty vector or the same vector with cloned *phoPQ* operon (pPQ). Middle bar = median; box limit = upper and lower quartile; extremes = minimal and maximal values. Disease index indicates: 0 no symptoms; 1-2 tissue damage mostly localised near inoculation site; 3-4 extensive tissue damage spreading sideways from the inoculation site. A Kruskal-Wallis H test showed a statistically significant difference in disease indices between the strains (p = 0.002). Three typical leaf sections for each variant are shown below the box plot. b) Potato tubers inoculated with JN42 wild type and *phoP* mutant bacteria (5·10^5^ or 1·10^5^ cells per inoculation site). Macerated tissue was weighed at 48 hpi.

### PhoPQ system controls over a hundred genes in *P. versatile*

RNA-seq transcriptional profiling revealed 116 genes with expression level differing between the wild type and the *phoP* mutant strains. Most of the differentially expressed genes (DEGs) were activated, only 28 were repressed. Table 3 and Supplementary table 1 depict the genes whose expression differed significantly (given a false discovery rate of 0.05) by at least 2-fold. The regulon composition and its control by PhoP are discussed in more detail in the following sections.

**Table 3.**
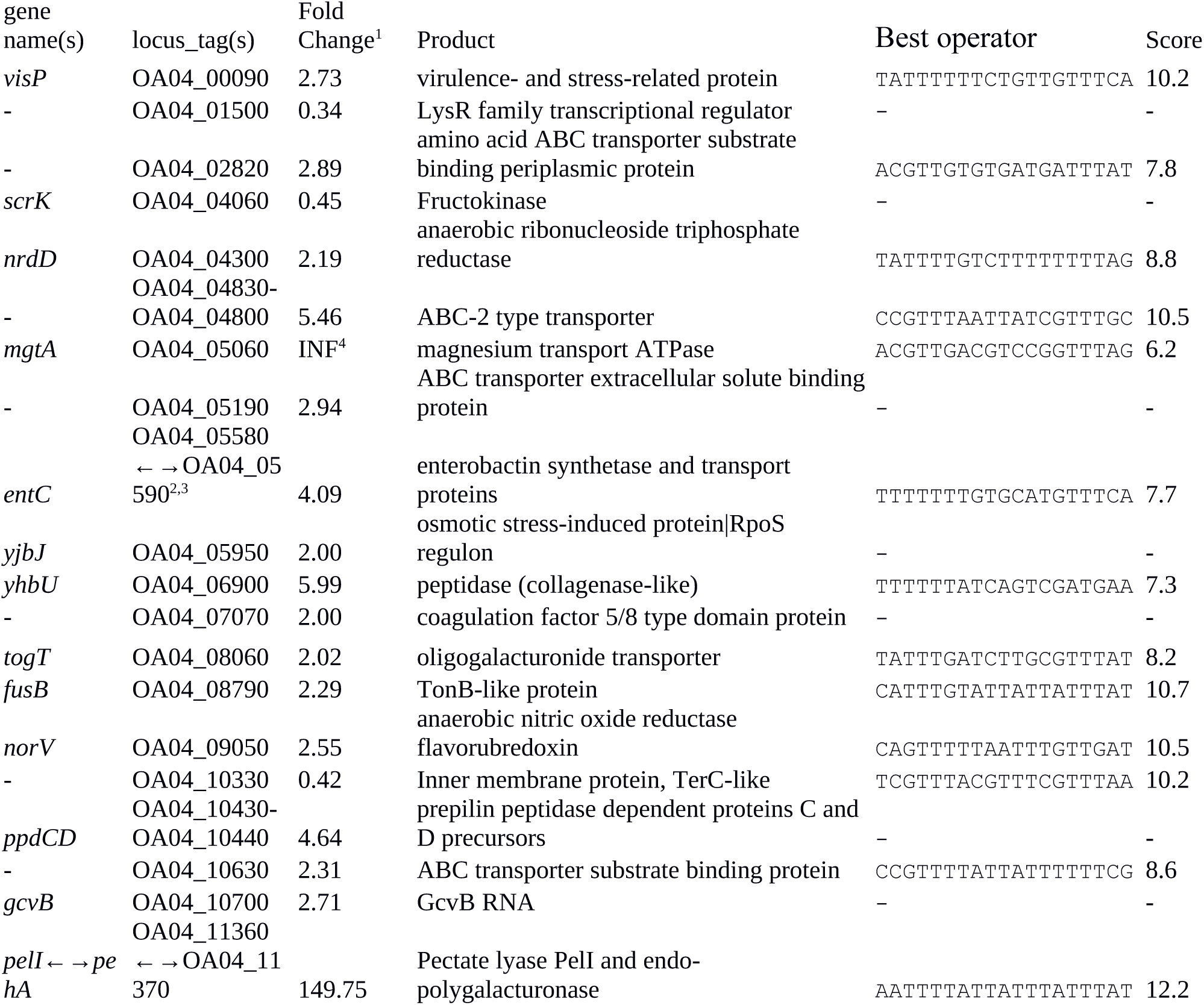
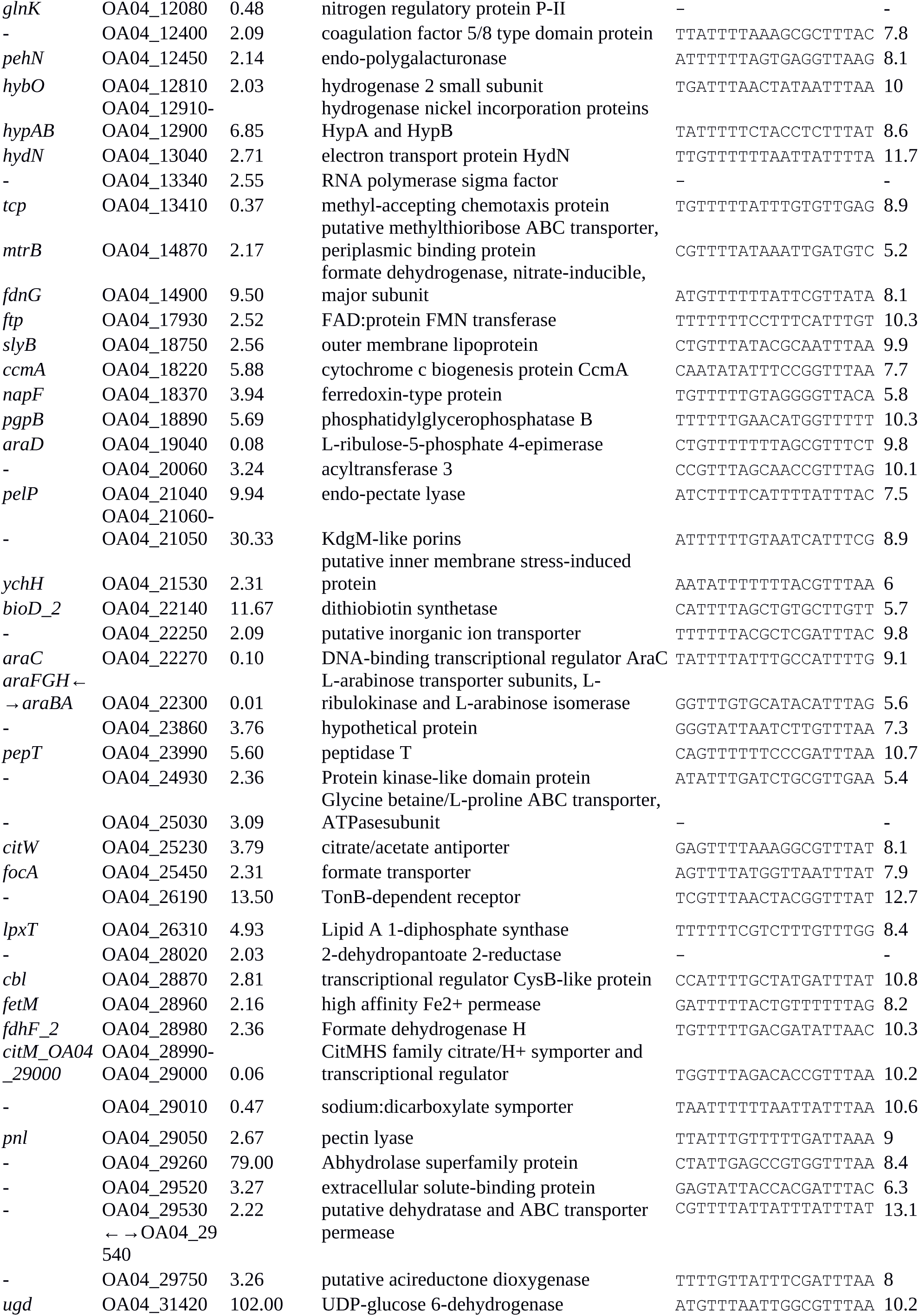
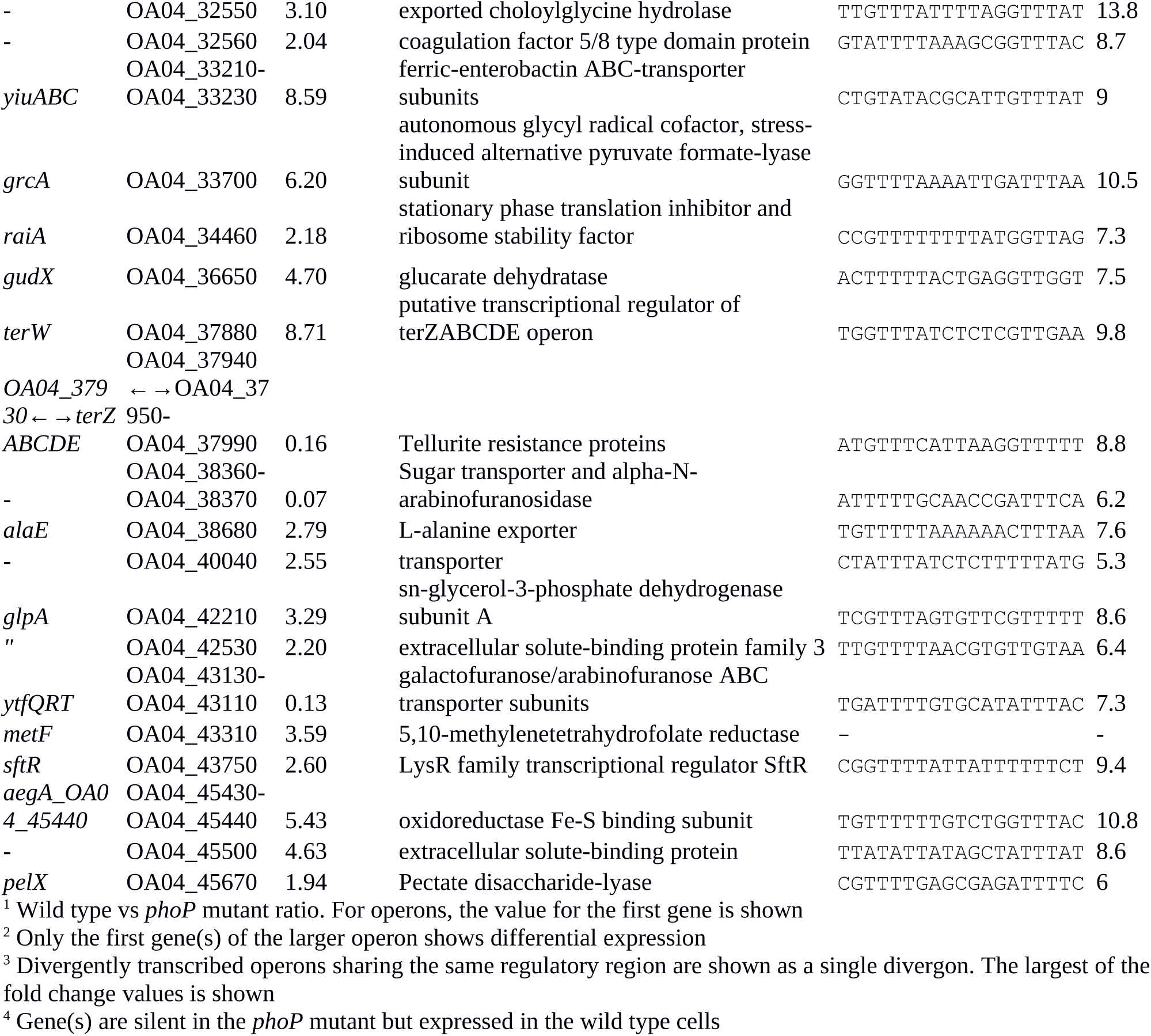
PhoP-dependent genes of *Pve*.

The RNA-seq data also confirmed the absence of *phoP* expression in the mutant cells. Expression of *phoQ* located immediately downstream in the same operon was barely detectable in the *phoP* mutant, which is probably due to the polar effect of *phoP* inactivation. Expression levels of the selected genes from different functional categories were verified by qPCR and found to correlate well with RNA-seq data (data not shown).

We looked for the presence of PhoP binding sites in the regulatory regions of the differentially expressed genes. ChIPMunk (39) could locate a relatively weakly conserved 19 bp sequence containing two direct gTTTa repeats (Figure 2A) in about half of the 85 regulatory regions of the differentially expressed genes. This sequence resembles the PhoP binding sites reported for other enterobacteria (Figure 2B and 2C) and was therefore considered the likely PhoP binding site.

**Figure 2.**
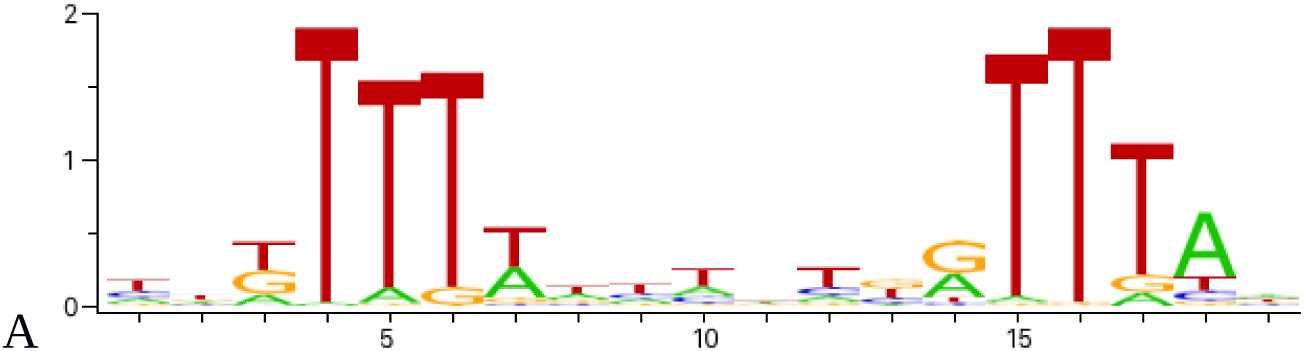
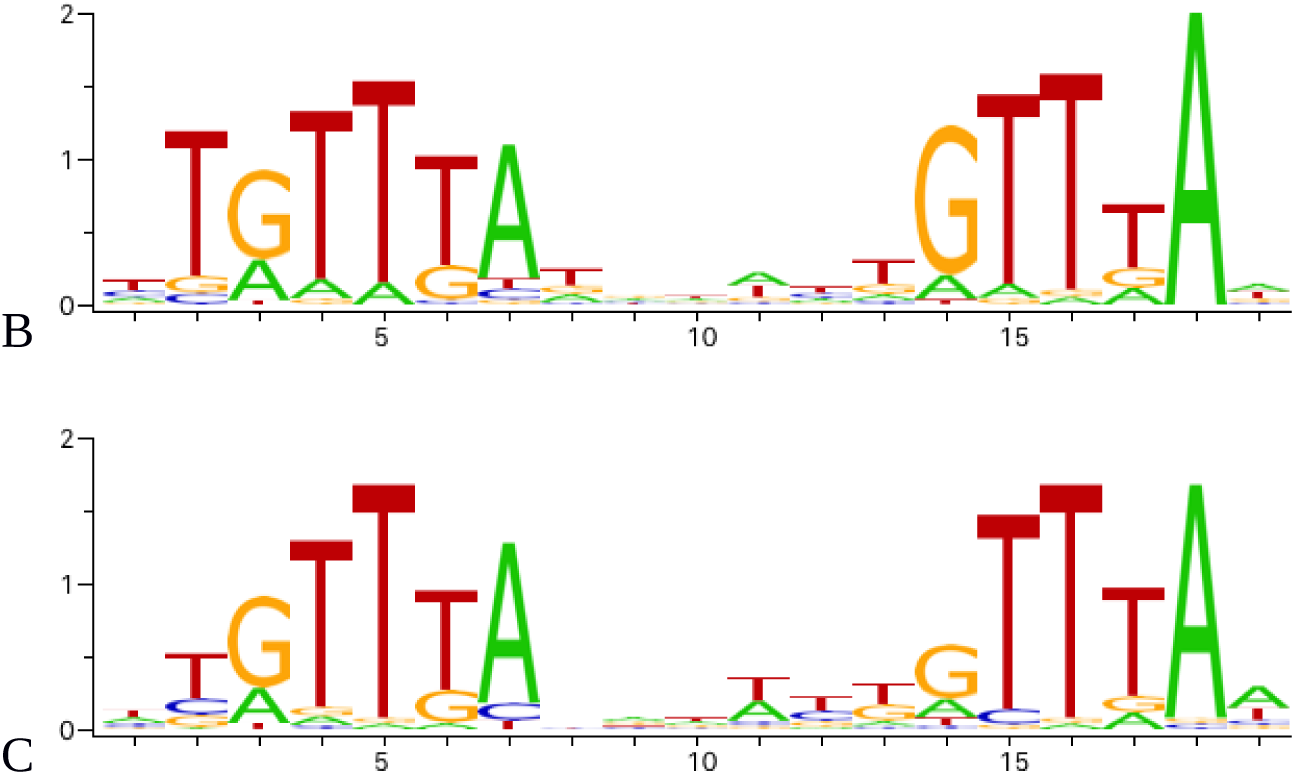
Sequence logos of operator motifs for three PhoP orthologues. A – *P. versatile* motif (made from the sites listed in Table 3). B – *E. coli* motif (combined non-redundant RegulonDB and CollecTF data). C – *S. enterica* motif (CollecTF data).

PhoP proteins from *Pve, E. coli* and *S. enterica* have about 80% identical residues across their full amino acid sequences. Analysis of their DNA binding domains shows identical amino acid residues in the positions making specific contacts with DNA bases (Figure 3). This strongly suggests that indicated proteins have very similar operator sequence specificities. The differences between their reported operator motifs may reflect the differences between the data sets and algorithms used for their inference. Therefore, we also used well known *E. coli* and *S. enterica* PhoP motifs to scan the *Pve* genome. This resulted in the identification of additional operators for the genes differentially expressed in this experiment.

**Figure 3.**
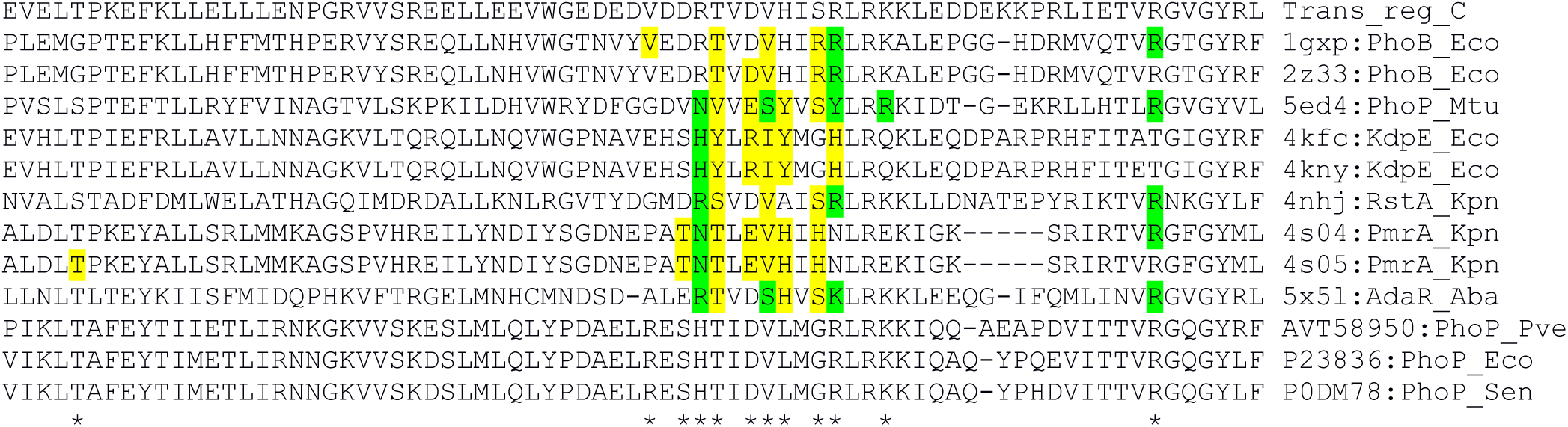
Alignment of DNA binding domain sequences of the OmpR family TFs. The sequences are aligned with hmmalign according to the Trans_reg_C (PF00486) family model. The amino acid residues of OmpR family proteins that form direct contacts with nucleotide bases were determined by the interaction service of NPIDB (43). The residues making specific contacts with DNA bases are highlighted in green (hydrogen bonds) and yellow (hydrophobic contacts). PDB/GenPept/Uniprot ID, protein name and species abbreviation are shown to the right of each sequence. Positions with contact observed in at least one sequence are marked with an asterisk.

### *The* P. versatile PhoP regulon part conserved in Enterobacterales

The universally conserved core of enterobacterial PhoP regulons is known to include only three genes: *phoP, phoQ* and *slyB* (25). In *Pve*, there are high-scoring PhoP-binding sites in front of both the *phoPQ* operon and the *slyB* gene. The expression level of *slyB* is decreased 2.6 times in the *phoP* mutant (Table 3). Therefore, these three genes are highly likely to be directly controlled by PhoP in *Pve*.

Additional PhoP regulon members shared by *P. versatile* with some other enterobacteria are required for cell envelope modifications that improve resistance to antimicrobial peptides and oxidative stress.

The eight genes of the *ugd-arn* locus are involved in the outer membrane lipopolysaccharide modification. The linked *ugd* gene and *arn* operon products add 4-aminoarabinose to the lipid A domain of the LPS, which reduces the negative charge and makes the cell more resistant to the cationic antimicrobial peptides (CAMPs) (45). These genes are expressed up to 100-fold stronger in the wild type *Pve* cells compared to the *phoP* mutant (Table 2). Seven *arn* genes are located downstream of the *ugd* gene and are transcribed in the same direction. A single gap (223 bp) within this locus is located between *ugd* and *arnB*. The gap contains a putative transcriptional terminator, indicating that *ugd* and *arnBCADTEF* might constitute separate transcriptional units. However, multiple high scoring PhoP binding sites are located in front of both *ugd* and *arnB*, ensuring direct PhoP control over the whole locus.

Expression of four more LPS-related genes is activated by PhoP 3-6 fold. The product of OA04_20060 ORF belongs to the acyltransferase 3 family (PF01757). Although the function of this ORF is not clear, at least some members of PF01757 (*S. enterica* OafA, *Staphylococcus aureus* OatA) are involved in the cell envelope modification by O-acetylating peptidoglycan (PG) which increases resistance to lysozyme (46,47). *visP* (*ygiW*) homologue from *S. typhimurium* codes for a virulence- and stress-related protein which binds to peptidoglycan and is required for polymyxin B resistance (48). *lpxT* codes for the lipid A 1-diphosphate synthase (49) which increases lipid A negative charge and therefore counteracts the action of *arn* gene products. *pgpB* codes for one of the three redundant phosphatidylglycerophosphatases which catalyse the dephosphorylation of phosphatidylglycerol phosphate to phosphatidylglycerol, an essential phospholipid of the inner and outer membranes (50). It also has undecaprenyl-pyrophosphate phosphatase activity, required for the biosynthesis of the lipid carrier undecaprenyl phosphate (51). Since LpxT uses undecaprenyl pyrophosphate as a phosphate donor to phosphorylate lipid A (49), PhoP-dependent PgpB activation might reduce phosphate donor availability for LpxT and hence prevent (or reduce) increase in negative charge of lipid A.

As the products of the *ugd-arn* locus and LpxT have the opposite effects on the lipid A negative charge and the CAMP resistance, we have checked resistance of *P. versatile* to cationic peptide polymyxin B and found it to be drastically reduced in the *phoP* mutant (Figure 4). Polymyxin resistance was restored in the phoP mutant to the wild type levels *by* a plasmid expressing the full phoPQ operon. Thus, the net phenotypic result of the PhoP-activated remodelling of the cell envelope in *Pve* is the activation of the CAMP resistance.

**Figure 4.**
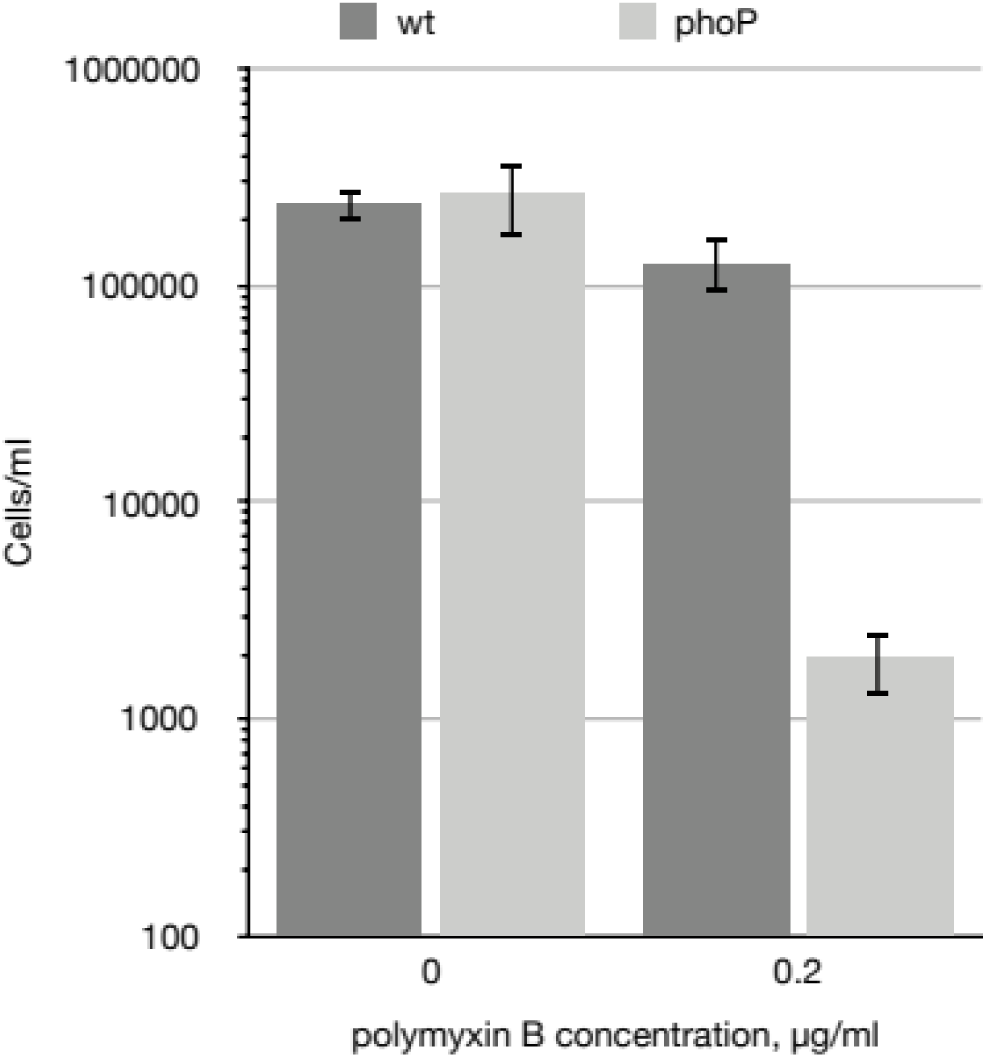
*phoP* is required for *Pve* polymyxin resistance. Wild type and *phoP* mutant *Pve* cultures were grown aerobically in LB medium to mid-log phase, diluted to 10^5^ cells ml^-1^ and split in two. 0.2 µg/ml of polymyxin B was added to one of the split cultures and incubation continued for 4 more hours. Surviving cell numbers were determined by plating. Mean values of three biological replicates and 95% confidence intervals are shown.

Several genes coding for stress-induced proteins, mostly poorly characterised, were found among differentially expressed. These include the following:

- *grcA* (6X activation) coding for the alternative pyruvate formate-lyase subunit,
- *raiA* (2.2x activation) coding for the stationary phase translation inhibitor and ribosome stability factor,
- *yjbJ* (2x activation) known to be induced by osmotic stress (52),
- *ychH* (2.3x activation) coding for an inner membrane protein induced by hydrogen peroxide and cadmium (53).

### Polygalacturonic acid utilization

PhoP was found to regulate a selection of pectinolysis related genes, activating eight and repressing one of them.

Just as in *P. parmentieri* (16), PhoP is required for *pehA* transcription in *Pve. pehA* expression demonstrates the strongest response to *phoP* inactivation (150x), which correlates with the presence of multiple PhoP binding sites in the *pehA* regulatory region.

Four more PhoP-activated genes code for pectinolytic enzymes: secreted endo-pectate lyase PelP (10x activation), periplasmic exopolygalacturonate lyase PelX (1.9x activation), secreted endopolygalacturonosidase PehN (2.1x activation) and pectin lyase Pnl (2.7x activation). Neither of these proteins is considered a major PGA depolymerase, although PelX was required in *D. dadantii* for fast growth on PGA (54,55).

Three oligogalacturonate transporter genes are also under positive PhoP control. The neighbour loci OA04_21050 and OA04_21060 (10x and 30x activation) code for porins with weak homology to oligogalacturonate-specific porins KdgM and KdgNand the *togT* gene (2x activation) codes for oligogalacturonate/cation symporter. The major oligogalacturonate transport and utilisation locus *kdgF kduDpelWtogMNAB kdgM* (1.5-1.7x activation) also has PhoP binding sites in the appropriate regulatory regions. The difference in expression levels suggests possible input of additional regulator(s) into transcriptional control of this locus.

Overall, PhoP appears to be a strong activator of extracellular and outer membrane components and a weak activator of periplasmic and cytoplasmic membrane components of polygalacturonate utilisation system. Together, PhoP-activated genes code for a complete set of proteins required for polygalacturonate depolymerisation and transport into the bacterial cytoplasm. However, *Pve* has more genes encoding PGA depolymerases that showed no PhoP dependence in our experiment. In particular, expression levels of the genes in the major pectate lyase gene cluster (*pelA, pelB, pelC, pelZ*) were the same in the wild type strain and the *phoP* mutant. It is therefore tempting to speculate that PhoP activates a subsystem of PGA depolymerisation and transport, finely tuned for a specific condition or a specific variant of PGA modification.

*pelI* is the only pectinolysis gene repressed (7x) by PhoP. It is located next to *pehA*, is transcribed divergently and shares its regulatory region with *pehA*. PelI is the major pectate lyase in planta (56). It was also reported to induce hypersensitive response in plants (57). Since high *pelI* expression can be expected to occur only when the PhoPQ system is inactivated, high *pelI* expression *in planta* may indicate such an inactivation.

### Citrate transport and utilisation

Two genes of citrate transport are controlled by PhoP: *citM* (17x repressed) and *citW* (6x activated).

*citW*, coding for citrate/acetate antiporter, is located within the citrate fermentation locus. The locus includes *citAB* operon encoding a citrate responsive two-component system and a divergently transcribed *citW*, which is followed by the *citYCDEFXG* operon (coding for the subunits of citrate lyase). All these genes (including *citW*) were poorly expressed in the conditions of our experiment.

*citW* and the whole citrate fermentation locus is expected to be relevant for citrate utilisation in the anaerobic conditions while *citM*, coding for a citrate/proton symporter, – in the aerobic ones (58,59). Therefore, PhoP can be considered as a switch between the respiratory and fermentation modes of citrate utilisation.

One more transporter gene, *OA04_29010*, is located downstream of the *citM* operon and codes for a putative sodium:dicarboxylate symporter. Just as the two preceding ORFs, *OA04_29010* is negatively controlled by PhoP (2.1x repression).

Besides the transporters and regulators, the *tcp* gene encoding citrate chemotaxis receptor is activated threefold in the *phoP* mutant.

### Arabinose transport and utilisation

14 genes related to transmembrane arabinose transport and utilisation are organised into three or four operons in *Pve*, negatively regulated by PhoP: the divergently transcribed *araFGHC* and *araBA* and the unlinked monocistronic *araD*. The *araC* gene is separated from *araH* by a 203 bp gap permitting an independent transcription initiation, but the lack of an obvious transcription terminator in this gap can also allow cotranscription with the upstream *araFGH* genes.

The *ytfQRTyjfF* operon encodes a periplasmic sugar-binding protein YtfQ and three subunits of an ABC transporter. YtfQ was shown to specifically bind the furanose forms of arabinose and galactose (58). Therefore, this operon is likely to code for an arabinofuranose or galactofuranose transporter (or a transporter of an arabinofuranose/galactofuranose containing oligosaccharide). The *OA04_38360*-*OA04_38370* operon codes for the putative glycoside-pentoside-hexuronide transporter and glycosyl hydrolase which belongs to the subfamily 43.26 according to CAZy classification (59). Bacterial members of this subfamily are annotated in CAZy as exo-α-1,5-L-arabinofuranosidases.

Arabinose is present in plant cell walls mainly in the form of arabinofuranosyl residues of the cell wall polysaccharides and proteoglycans such as pectic arabinan, arabinoxylan, and arabinogalactan-proteins (60). Presumably, the *OA04_38360*-*OA04_38370* and *ytfQRTyjfF* operon products are involved in transport and utilisation of arabinofuranose containing products of plant cell wall breakdown. As the *OA04_38370* product does not have a signal peptide, it can only degrade cytoplasmic substrates like an α-L-Ara*f* containing oligo- or disaccharides which could be released from plant cell wall by the action of other enzymes. At least two enzymes of *Pve* 3-2 genome are relevant: a putative arabinogalactanase, encoded by *OA04_08560*, and an arabinogalactan endo-1,4-beta-galactosidase, encoded by *ganA* of the *ganEFGAB* operon. The *ganEFGAB* operon is controlled by the GalR, as well as the *OA04_38360*-*OA04_38370* operon. Although no differential expression was registered *in vitro* for any of the two loci encoding secreted arabinogalactanases, both arabinogalactanase genes could be expected to be induced by galactose *in planta* via GalR. Indeed, our recent transcriptome profiling of *P. atrosepticum* has shown 4- and 24-fold *in planta* induction of *OA04_08560* and *ganA* orthologues *ECA0852* and *ECA3128* (27).

High scoring PhoP binding sites are located near two out of six arabinose related differentially expressed transcriptional units. A PhoP binding site overlaps the very beginning of the *araD* reading frame, suggesting direct negative control of *araD* by PhoP. Another binding site was found in front of the *araC* gene in a position fit for repression. Three rather weak PhoP binding sites were found in different ChIPmunk runs within the regulatory region of the *OA04_38360*-*OA04_38370* operon, suggesting the likelihood of direct PhoP control over this operon as well. Indirect control seems mote likely for the remaining three arabinose-related transcriptional units.

Since *araC* codes for the arabinose-responsive transcription factor, we searched for AraC binding sites in the *Pve* genome. First, the AraC operator profile was constructed. For this, we downloaded the AraC_Enterobacterales motif from RegPrecise (61). However, this short (17 bp) motif corresponds to just a part of the binding site. AraC has two DNA binding domains, both can contact DNA and the active AraC form is dimeric (62), so the real binding site should be much wider. We checked the conservation around the sites annotated in RegPrecise and found the conserved region to cover about 40 bp – sufficient to allow binding of four DNA binding domains of the dimeric AraC. The extended part of the motif is less conserved, but has an overall resemblance to the conserved part and has the same orientation (Figure 5). This configuration of the operator is suitable for the binding of a tandem of AraC subunits in their activating conformation. The most conserved part of this motif corresponds well to the one recently defined experimentally (63).

**Figure 5.**
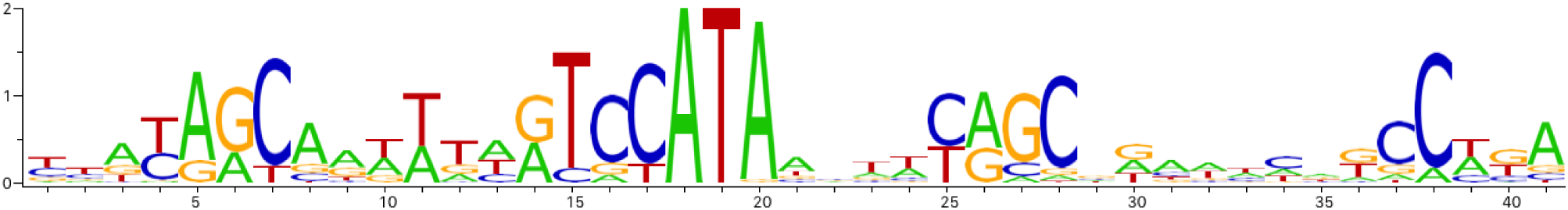
AraC binding site sequence logo.

Scanning of the *Pve* genome with the AraC profile has found matches in three locations: in the regulatory region between the *araBA* and *araFGHC* operons (two sites), in front of the *araD* gene and in front of the *ytfQRTyjfF* operon. Therefore, PhoP dependence of differentially expressed arabinose-related transcriptional units could be explained by direct negative control by PhoP over *araD, araC, ytfQRTyjfF* and *OA04_38360*-*OA04_38370*, combined with positive control via AraC over *ytfQRTyjfF* and the other three operons that lack PhoP sites in their regulatory regions.

Further investigation of PhoP-dependent control over arabinose utilisation genes has shown, that (i) arabinose is a strong inducer of *ara* genes and (ii) the observed differential expression depends on the presence of sodium polypectate in the medium (Figure 6). Without polypectate, no PhoP dependence of *ara* genes is observed, both with and without arabinose. Sodium polypectate appears to be a weak inducer of arabinose-related operons in the wild type strain, but induction is much stronger in the *phoP* mutant (Figure 6). PhoPQ two-component system does not seem to respond to polypectate, as most of the other DEGs have the same expression with and without polypectate (data not shown). This suggests indirect control by another PhoP-controlled and polypectate responsive transcriptional regulator and requires further investigation.

**Figure 6.**
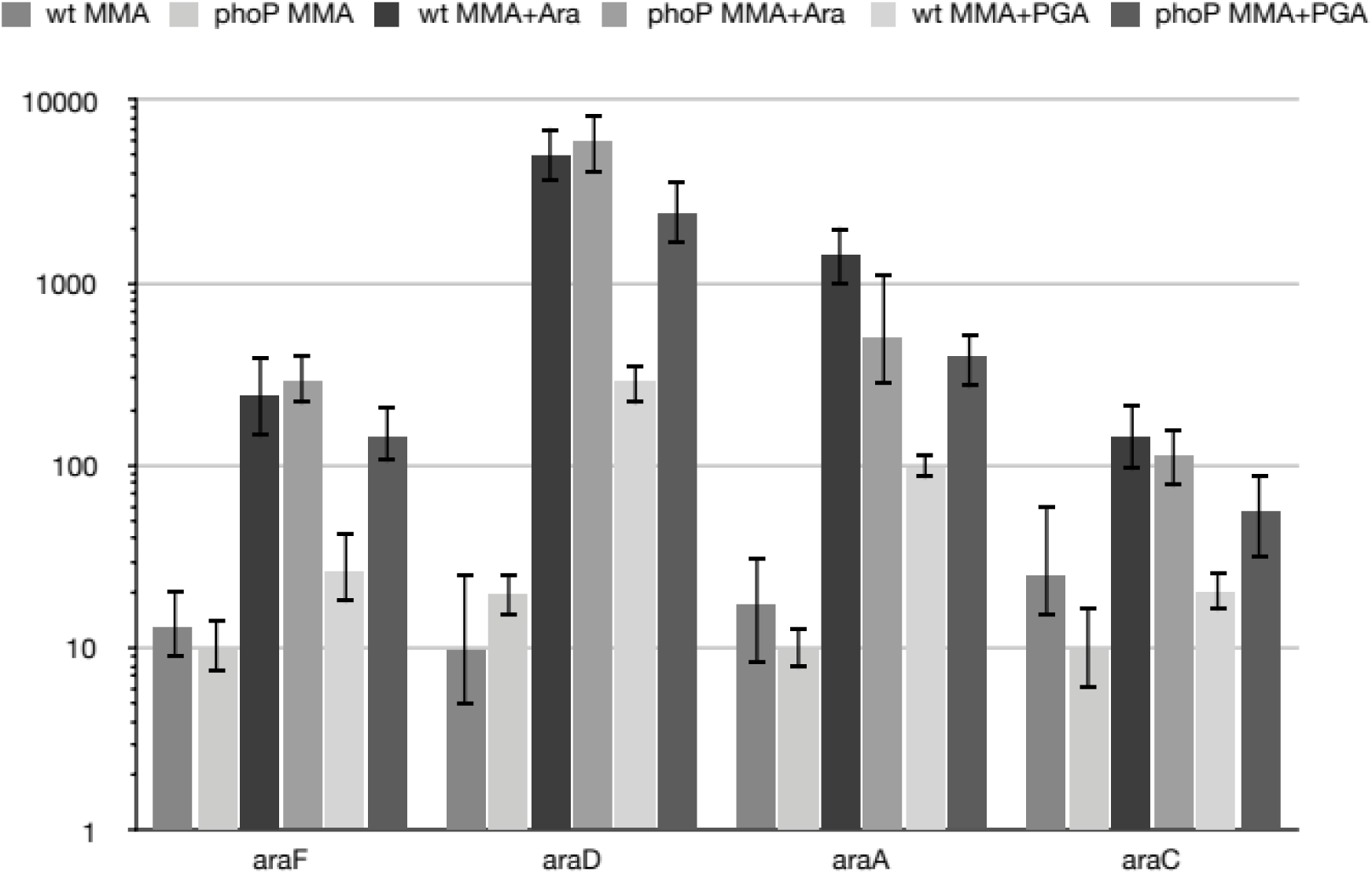
Differential expression of *ara* genes depends on the presence of polygalacturonate For qPCR measurement, RNA was isolated from the wild type (wt) or *phoP* mutant *Pve* cells grown in MMA, MMA with 0.5% arabinose (MMA+ara) or in MMA with 0.5% sodium polypectate (MMA+PGA).

### Other transporters

PhoP is known to be involved in the regulation of at least one of the Mg^2+^ transporters in enteric bacteria (25). *P. versatile* seems to have the only dedicated Mg^2+^ transporter encoded by *mgtA*. In the conditions of our RNA-seq experiment with moderate (0.5 mM) Mg^2+^ concentration *mgtA* was weakly expressed in the wild type cells and was completely silent in the *phoP* mutant. At least tenfold higher expression of the 5’ UTS compared to the coding frame of *mgtA* (Supplementary figure 3) hints to a riboswitch-like control of transcription elongation into the coding region of *mgtA* analogous to the mechanism described for *S. enterica* (64).

PhoP activates several genes and operons involved in iron acquisition. Since our experiment was performed without iron limitation, these genes had a rather low expression and differential expression in many cases was detected only for the first gene of multigene operons. These genes and operons include *fetM* encoding high-affinity Fe^2+^ permease, *yiuABC* coding for ferric-enterobactin ABC transporter and OA04_26190 encoding TonB-dependent outer membrane siderophore receptor. *entC*, the first gene of the enterobactin-like siderophore synthesis operon *entCEBFA*. Putative permease gene *OA04_05580* preceding *entC* and divergently transcribed is also PhoP activated and might be involved in siderophore transport. *fusB*, the first gene of the *fusBACD* operon, encodes a TonB-like protein known to be required for import of plant ferredoxin (65) via the outer membrane ferredoxin receptor FusA (66). Another putative transporter locus OA04_13300-OA04_13380 encodes all components of cell surface signalling system including TonB-dependent ferrichrome-iron receptor, TonB-, ExbB- and ExbD-like proteins, FecI-like sigma-factor (OA04_13340) and FecR-like anti-sigma factor. Additional components encoded here include an M16 family protease distantly related to FusC (OA04_13370) and FusD-like ABC transporter (OA04_13360). This arrangement allows us to speculate about the involvement of this locus in the acquisition of iron, possibly complexed with a protein carrier. The FecI-like sigma factor gene *OA04_13340* is expressed better in the presence of PhoP, and three putative promoters controlled by OA04_13340 (Supplementary figure 4) could be located in front of transcriptional units of this locus.

Three PhoP regulated transporters participate in amino acid import. The *OA04_29540-OA04_29570* operon is likely involved in the import and modification of branched-chain amino acids. *OA04_42530* codes for the periplasmic binding protein of another amino acid transporter. OA04_02820 product is annotated as a cystine-binding periplasmic protein.

Three more controlled loci encode putative efflux transporters. These include L-alanine exporter AlaE (OA04_38680), putative multidrug efflux transporter OA04_04830-OA04_04800 and AraJ family efflux permease OA04_40040.

Three additional transporters do not belong to any of the groups mentioned above. *mtrB* codes for the periplasmic binding subunit of methylthioribose ABC transporter. *OA04_25030* encodes the ATPase subunit of glycine betaine/L-proline ABC transporter, *OA04_22250* – the putative sodium/ bile acid cotransporter. The last transporter may act in conjunction with choloylglycine hydrolase encoded by *OA04_32550*, which is also activated by PhoP.

The potential substrates of four other transporters (OA04_10630, OA04_29520, OA04_45500 and OA04_05190-OA04_05160) could not be inferred even approximately.

All the genes coding for the transporter subunits described above are activated by PhoP. However, in many cases the observed effect of *phoP* inactivation may be indirect. In particular, we found no PhoP binding sites near *mtrB, OA04_25030, OA04_05190* and only weak sites could be located for *OA04_38680, OA04_45500* and *OA04_02820*.

We also note that the GcvB ncRNA, a well-known negative regulator of amino acid transporter genes (67), is 2.7x activated. This effect may partially compensate for the decrease of the transporter gene expression in the *phoP* mutant. We could not find a PhoP binding site in front of *gcvB*. Therefore, *gcvB* expression control by PhoP might be indirect, just as has been reported for *E. coli* (68). GcvB has also been reported to control PhoP expression posttranscriptionally by both destabilising its mRNA and decreasing its translation (69). Such reciprocal repression between PhoP and GcvB, if present in *Pve*, might result in amplification of any changes in expression of either regulator.

### Tellurite resistance genes

A group of seven genes (*terZABCDE* and divergently transcribed *OA04_37930*) tentatively annotated as coding for tellurite resistance proteins is 2-6x repressed by PhoP. *OA04_37930* is the first gene in the operon (1.7-1.9x repression of the four downstream genes). These genes are a part of a large locus of 18 presumably functionally linked genes (70) organised into four or five operons. Although the exact function of this locus *in vivo* is not clear, orthologous genes are required for tellurite resistance in some bacteria (71–73). Tellurite is also thought to induce oxidative stress and tellurite resistance might be the result of decreased sensitivity to the oxidative stress (74).

The unlinked *OA04_10330* locus (2.4x repression, codes for a TerC-like membrane protein) has a “good” PhoP binding site, suggesting direct PhoP control in *Pve*. At least one PhoP binding site is located in the regulatory region between *terZ* and *OA04_37930*, one more is present upstream of the *terW. terW* (8.7x activation) is located a few genes away and was shown to be responsible for transcriptional control of ter genes in *E. coli* (77). The region most conserved between *E. coli* and *Pectobacterium sp*. sequences contains a 25 bp palindrome (Figure S2).

*Pve* 3-2 cells growing on the medium with low potassium tellurite concentration form characteristic black colonies, indicating the reduction of tellurite to tellurium. We did not notice a significant difference in black colouring between the wild type *Pve* and the *phoP* mutant (data not shown), but the *phoP* mutant could grow in LB medium with tellurite concentration 3 µg ml^-1^ which is inhibitory to the wild type strain (data not shown).

In *E. coli*, its position corresponds to the TerW-regulated *terZABCDE* supposed promoter region (77). In *Pve*, this palindrome is located in the middle of the intergenic region between the divergently transcribed operons *(terZABCDE* and *OA04_37930-OA04_37890*), therefore TerW could control both of them. Thus, either the direct repression or a transcriptional cascade, where PhoP activates *terW* and TerW represses divergent operons, are supposed to explain the observed PhoP dependence of the genes in the *ter* locus.

The actual role of “tellurite resistance” genes in a soft rot pathogen requires separate investigation. Soft rot bacteria can hardly ever encounter significant concentrations of tellurite, so these genes are likely responsible for coping with some other stress. Among *Pectobacteriaceae*, “tellurite resistance” genes are only present in three pectobacteria: *P. versatile, P. parmentieri* and *P. polaris* suggesting such stress is important for the lifestyles of only these three species.

### Anaerobiosis-related genes

The transcription profiling experiment described here was performed in aerobic conditions, and most of the anaerobiosis-related genes were expressed poorly. However, the PhoP-dependent positive control over important aspects of anaerobic metabolism was noticeable. The largest group of PhoP-activated genes is required for the detoxification of formate produced by enteric bacteria in anaerobic fermentative conditions. These genes include the formate transporter gene *focA* and several subunits of formate hydrogenlyase. Although the *hyf* operon encoding the hydrogenase 4 was silent in the conditions of our experiment, other weakly expressed operons within the hydrogen metabolism gene cluster were PhoP dependent. These include three operons. The first operon is *hybOABCDE*, coding for subunits of hydrogenase 2. The second one is a *hypABCDE* operon coding for hydrogenase maturation proteins. The third operon includes genes coding for electron transport protein, which is required for formate dehydrogenase activity (*hydN*), formate dehydrogenase H (*fdhF*) and one more hydrogenase maturase (*hypF*). The *Pve* 3-2 genome carries a second *fdhF* paralogue (*fdhF_2*) which is also PhoP activated. The *fdnGHI* operon, coding for nitrate inducible formate dehydrogenase N, is PhoP activated too.

Expression of the *aegA* operon, coding for formate metabolism proteins, is PhoP dependent. *aegA* codes for an oxidoreductase that was recently shown to be involved in formate-dependent catabolism of urate (75). A gene downstream *aegA, OA04_45440*, codes for an uncharacterised [4Fe-4S] ferredoxin subunit of hydrogenase or formate dehydrogenase.

Anaerobic nitrate reduction may also be controlled by PhoP. The targets include a PhoP-activated *nap* operon responsible for periplasmic nitrate reductase production (differential expression was registered for only the first gene *napF*). Clear differential expression was detected for *ccmA*, the first gene of the *ccmABCD* operon coding for the subunits of the heme trafficking system. This system delivers heme to holocytochrome *c* synthase CcmFGH. And since this cytochrome *c* biogenesis system is required for maturation of cytochrome *c* subunits of nitrate and nitrite reductases, PhoP might be required for anaerobic nitrate and nitrite reduction. An additional layer of PhoP control over nitrite reduction is provided via regulation of formate dehydrogenase, which serves as the electron donor for formate-dependent nitrite reductase coded by *nrf* operon (poorly expressed in this experiment). PhoP also appears to control the *norVnorW* operon coding for anaerobic nitric oxide reductase.

Anaerobic respiration with fumarate as an electron acceptor may also be PhoP dependent due to control of at least one key gene, *glpA*. The *glpABC* operon codes for subunits of anaerobic glycerol-3-phosphate dehydrogenase, which is required for anaerobic respiration with fumarate as a terminal electron acceptor.

The *nrdD* gene codes for anaerobic ribonucleoside triphosphate reductase, which is essential in *E. coli* for deoxyribonucleotide synthesis during strict anaerobic growth (76).

### Regulators

Nine regulatory genes showed differential expression in our experiments. Three of the transcription factors (AraC, TerW and OA04_13340) and the regulatory RNA GcvB were already discussed above. *glnK* codes for the second nitrogen regulatory protein P-II. Since it is functionally equivalent to P-II encoded by *glnB*, and *glnB* expression level is much higher, the significance of the twofold difference of *glnK* expression levels should be minor.

OA04_01500 is a member of the RpoN regulon, had very low expression in the nitrogen-rich conditions of this experiment and was therefore unlikely to contribute significantly to the observed expression values.

Three genes, coding for LysR family TRs, are PhoP-controlled: *sftR* (2,6x activated), its close paralog *OA04_28450* (below the twofold threshold activated, statistically significant) and c*bl* (2,8x activated). The two paralogues are located within two unlinked clusters presumably involved in alkanesulfonates utilisation. We located the only potential operator in *Pve* genome, that might be bound by these two TFs in the atsR-atsBCA intergenic region (data not shown). Therefore, SftR and OA04_28450 involvement in the control of the DEGs identified in this experiment is unlikely.

*cbl* codes for a CysB-like protein, which orthologue in *E. coli* controls genes required for aliphatic sulfonate and taurine utilisation and homeostatic response to sulfate starvation, according to RegulonDB. The DBDs of orthologues show high similarity, unlike the ligand-binding domains, which suggests a possibility of binding different ligand(s) by these two TFs. CysB-like proteins were reported to control functions unrelated to sulfur utilisation (80–82). A rather high transcription level of *cbl* in *Pve* supposes this TF control of some DEGs in this experiment.

One more well-expressed transcription factor belongs to the GntR family. It is coded for by the *OA04_29000* gene located downstream of *citM*, probably in the same operon. The involvement of *OA04_29000* and *cbl* gene products in transcriptional control of the DEGs described here is currently under investigation and will be reported separately.

### PhoP regulon is activated in *Pve* by Ca^2+^ and Mg^2+^ limitation

In an attempt to find the conditions that might be responsible for PhoPQ activation in *Pve*, we checked some environmental stimuli known to be detected by the PhoQ sensor. There was no reaction to cationic peptide polymyxin and low pH (data not shown), but the response to divalent cations was significant. Expression levels of PhoP-activated genes were the highest at low (10 µM) concentrations of both Ca^2+^ and Mg^2+^ (Figure 8). Increasing either cation concentration to 10 mM reduced expression levels of PhoP-activated genes. Importantly, expression of *phoP* was decreased 13 times by 10 mM Mg^2+^ and 24 times by 10 mM Ca^2+^. Expression level differences of *pehA* and *ugd* caused by divalent cation addition were in the range 16-35 times. Comparison with the *phoP* mutant strain grown in the same conditions shows that some active PhoP must be present in the wild type cells even after divalent cation addition as *phoP* inactivation results in larger (91-353 times) drop of *pehA* and *ugd* expression. We interpret these results as both Mg^2+^ and Ca^2+^ can bind PhoQ and significantly reduce its kinase activity required for phosphorylation of PhoP which strongly reduces transcription initiation at PhoP-activated promoters.

**Figure 7.**
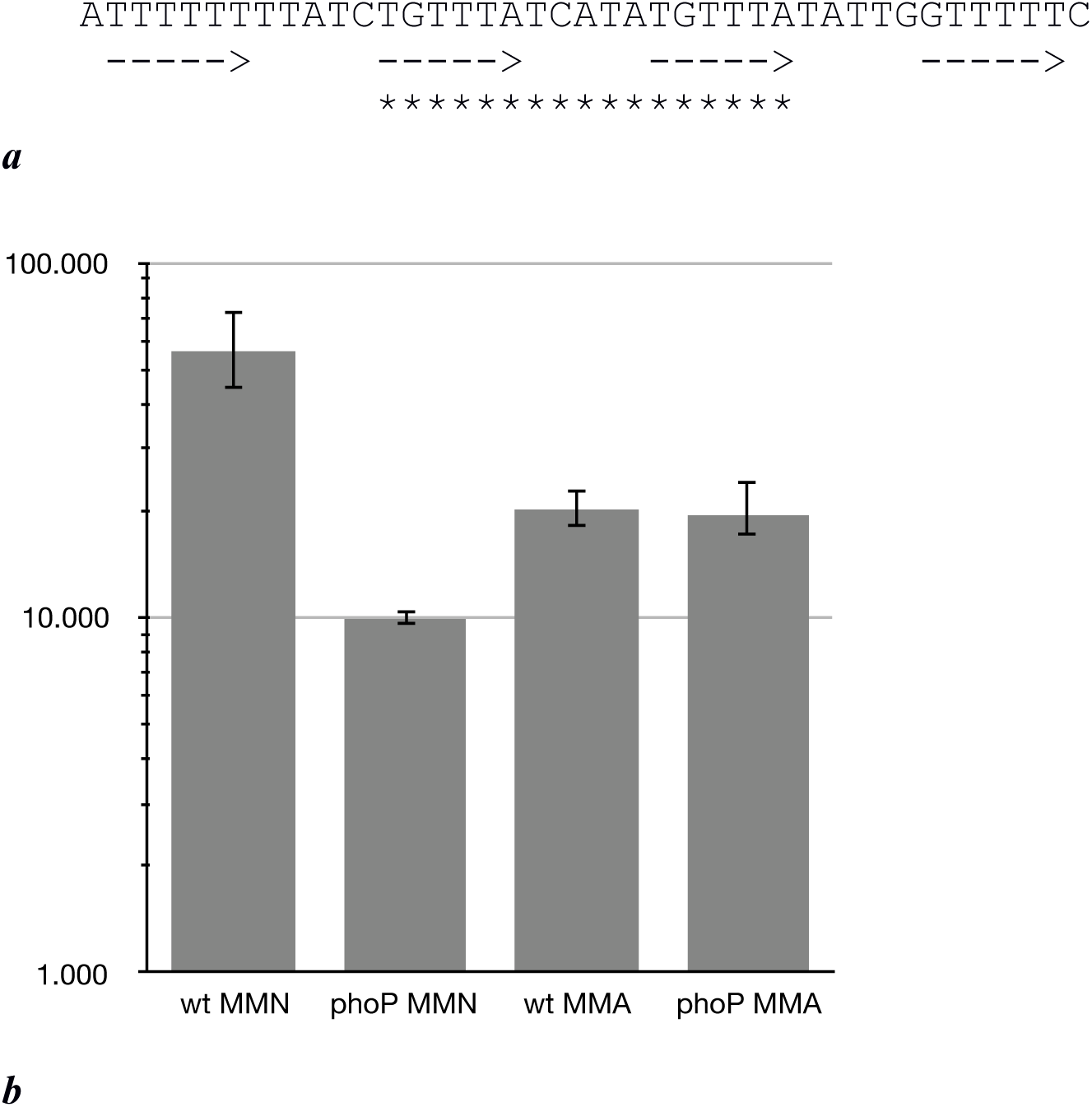
PhoP binding sites in the *expI* promoter region. a – The sequence of the PhoP binding sites in *expI* promoter region. Arrows indicate possible positions of PhoP monomer binding sites, asterisks – the most likely position for binding of PhoP dimer. b – *expI* expression in *Pve* cultures grown to 10^8^ cells ml^-1^ in MMN (10 µM Ca^2+^, 10 µM Mg^2+^) and MMA (10 µM Ca^2+^, 0.5 mM Mg^2+^).

**Figure 8.**
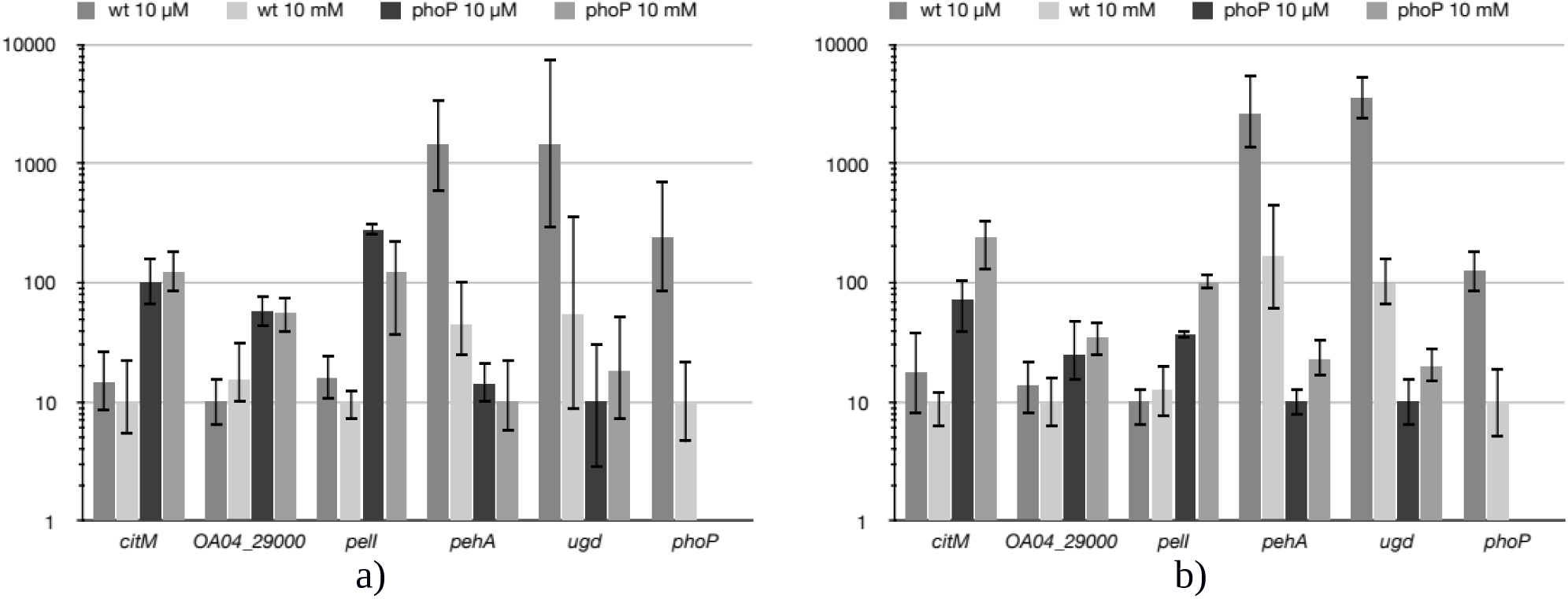
Ca^2+^ (a) and Mg^2+^ (b) reduce expression of PhoP-activated genes, but don’t affect PhoP-repressed ones.

Unexpectedly, divalent cation concentrations had little effect on PhoP-repressed genes. Expression levels of the three genes tested (*citM, pelI* and *OA04_29000*) differed less than two-fold in the wild type strain and these differences were not statistically significant (Figure 8). We conclude that (i) PhoP amounts present in the wild type *Pve* cells grown at 10 mM Ca^2+^ or Mg^2+^ are sufficient to cause repression of the two transcriptional units involved and (ii) phosphorylation may not be required for PhoP to bind its operator sites. The ability of reduced PhoP amounts to repress transcription of *pelI* and the *citM-OA04_29000* operon is not surprising, as regulatory regions of both transcriptional units have multiple high scoring (and presumably high affinity) PhoP binding sites.

### The extended PhoP regulon

Since PhoP is a global transcriptional regulator, PhoP-dependent genes differentially expressed in the single condition studied in this work probably constitute just a subset of the PhoP regulon. To see a broader picture, we analysed potential PhoP operators located near transcriptional units that did not pass differential expression thresholds. This analysis was done by scanning *Pve* genome sequence with four PhoP operator profiles: the ones based on our experimental data, *E. coli* data, *S. enterica* data and *in silico* inferred profile (Supplementary file 1). The scan identified a total of 812 genes organised into 254 transcriptional units that could be the targets of transcriptional control by PhoP (Supplementary file 2) and some can be genuine PhoP regulon members. Many of these putative PhoP regulon members are related to plant cell wall degradation and utilisation of carbon sources abundant in plants. Of note are the genes/operons encoding pectate lyases (*pelA* and *pelB*), pectin methyl/acetyl esterases (*pmeB* and *paeX*), three beta-glucoside transport/utilisation operons (*bglHDJ, bglYK arbFBH*) and five di/tricarboxylate transporters (*dcuA, dcuB, dctA, maeN* and *yflS*). For most of these genes, the PhoP binding site suggests an activator position. At least one negative regulatory site (that could block PhoP-dependent transcription activation) could be located between PhoP operator and translation start site (attenuators in front of *bglH, bglY, arbF* and possibly *paeX*, operators for DcuR in front of *dcuA, yflS* and *dcuB*, operators for NarP in front of *dcuB* and *yflS*).

A “strong” PhoP binding site is located in front of the *expI* gene coding for acyl-homoserine lactone synthase (a key regulator of quorum sensing). ExpI dramatically effects on gene expression (83), including virulence genes, therefore PhoP control of *expI* transcription was studied in more detail. The regulatory region of *expI* (Figure 7a) has four equally spaced sites (two central ones perfectly match the gTTTA consensus) for the binding of two PhoP dimers (Figure 7b). Since PhoP homologues can form oligomers (77), binding of a PhoP tetramer is also possible in this region.

The RNA-seq data showed a small (22%) difference in *expI* expression levels between the wild type and the mutant strains. The difference increased up to five-fold if the bacteria were grown to slightly lower cell density in the MMN medium at 10 µM concentrations of calcium and magnesium ions when the PhoPQ system is fully induced (Figure 7*c*).

### PhoP regulon expression *in planta*

Due to autoregulation, PhoP involvement in the adaptation to the plant environment must be accompanied by changes in its expression level. We have previously noticed decreased *phoP* expression in another example of *Pectobacterium*-plant interaction. A 3.5- and 5.2-fold *phoP* repression was observed for *P. atrosepticum* strain SCRI 1043 in asymptomatic and necrotic zones of infected tobacco stems (27). This reduction of *phoP* expression correlated well with expression level changes of PhoP controlled genes: e.g. *pehA* expression was reduced while *pelI* and *ara* genes expression was increased.

In the current investigation, *phoP* expression decreased about 4 times in macerated potato tuber tissues (Figure 9). In the bacterial cells, isolated from the rotten tuber tissues, expression level differences of PhoP regulon members between the wild type and mutant strains were much less pronounced than *in vitro* or even absent (Figure 9). This could be attributed to the partial inactivation of the PhoPQ system *in planta* as discussed in the following section. A single notable exception was the *pelI* gene, which had similar differences of expression levels between the wild type and *phoP* mutant strains *in planta* and *in vitro*. This could be attributed to a large number of high affinity PhoP binding sites in front of *pelI*. Since both dimerisation and DNA binding of PhoP is known to be independent on its phosphorylation (78), moderately reduced amounts of unphosphorylated PhoP that should be present in the cells from rotten tissues could be sufficient to cause *pelI* repression.

**Figure 9.**
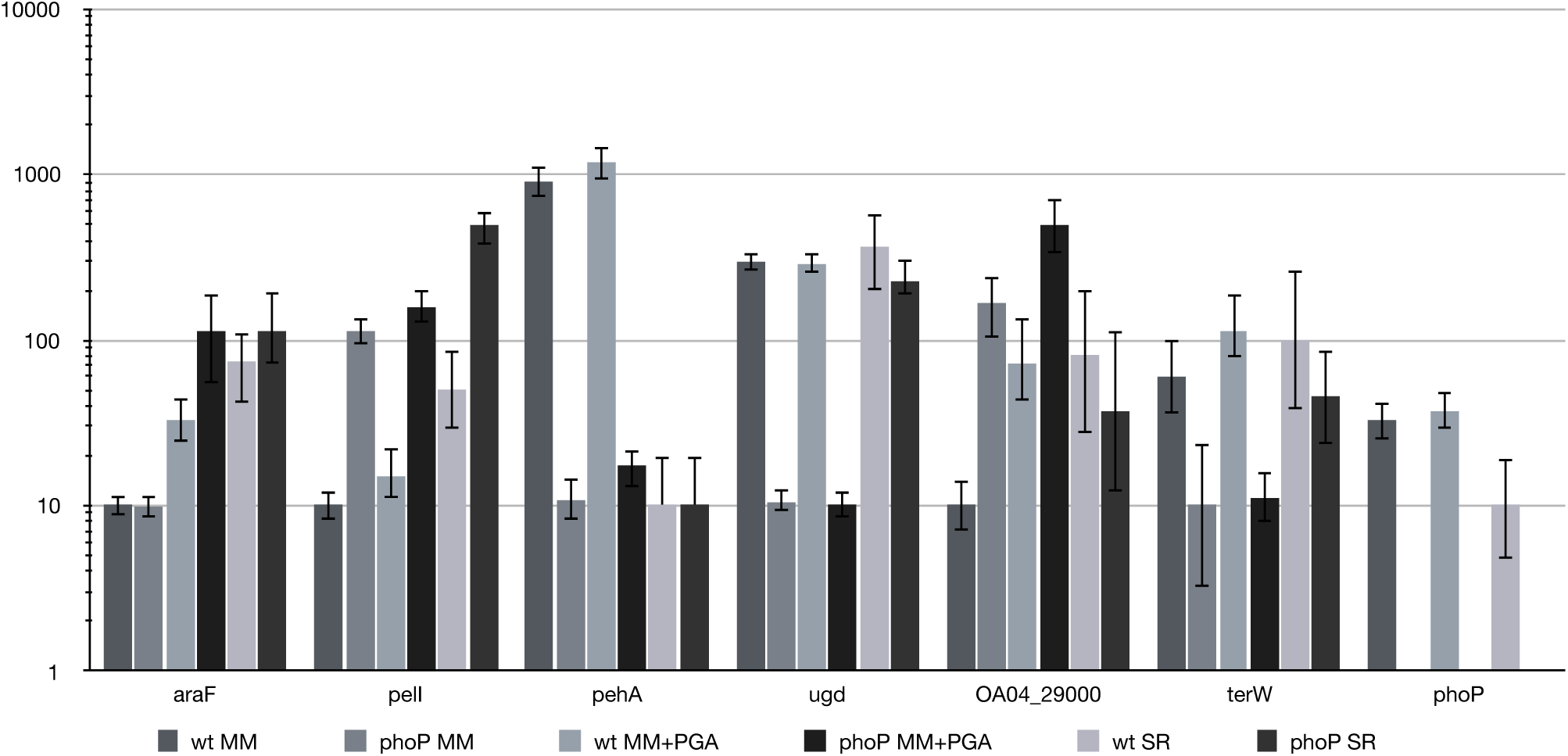
Relative expression levels of PhoP regulon members *in vitro* and *in planta*. For qPCR measurement, RNA was isolated from the wild type (wt) or *phoP* mutant *Pve* cells grown in minimal medium (MM), minimal medium with 0.5% sodium polypectate (MM+PGA) or from the cells of rotten potato tissues (SR).

## Conclusions

Our experimental results and *in silico* analysis show, that multiple additional transcription units are placed in *Pve* under the control of the highly conserved enterobacterial two-component PhoPQ system. The majority of the genes added to the PhoP regulon in *Pve* codes for the functions with the obvious importance for a soft rot plant pathogen (Figure 10). As a result, the PhoPQ two-component system can be considered a major modulator of *Pve* gene expression during the infection process. From the very early moments till the latest stages of infection, PhoP is involved in adjusting the expression levels of the major virulence factors, stress resistance factors, transporters and enzymes necessary for the utilisation of plant-derived products. This adjustment is possible due to the decrease of *phoPQ* expression at later stages of infection, likely caused by an increase in the concentrations of free divalent cations (both Ca^2+^ and Mg^2+^). Ca^2+^ was shown to be mostly cell wall-bound but is released when the cell wall is degraded by pectobacteria (79). Mg^2+^ concentration in plant apoplast was also reported to be quite low (80). At the onset of infection, PhoPQ expression is high, resulting in the activation of the *pehA* gene expression. High polygalacturonase activity at the first stage of infection may change the availability of other cell wall components and would release cell wall-bound Ca^2+^ and Mg^2+^, leading to the repression of *phoPQ* transcription and activation of PhoP-repressed genes, including *pelI*.

**Figure 10.**
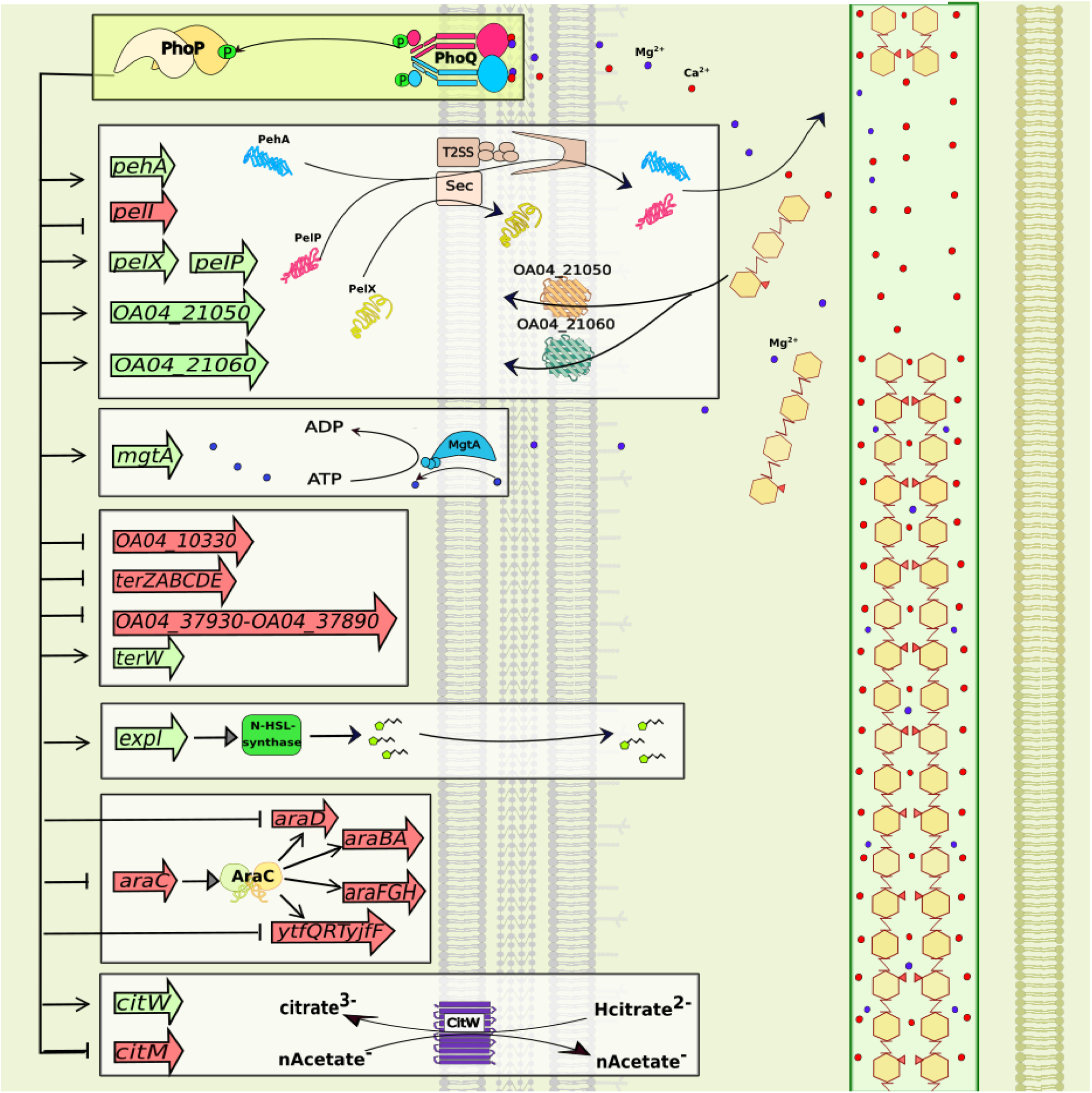
PhoPQ – dependent switch of virulence gene expression in *Pve*.

Only the genes with the obvious virulence-related functions are shown. The picture captures the active state of the PhoPQ system (low calcium/magnesium ion concentration), therefore the products are only shown for PhoPQ-activated genes. Activation/repression is shown as pointed/blunt arrows and activated/repressed genes are shaded green/red.

PelI is highly active in the presence of high Ca^2+^ concentrations, is known to be highly expressed *in planta* and is thought to be especially effective at the degradation of the calcium-rich middle lamella (81,82). High *pelI* expression, observed in potato tubers at late infection stages, may have a major effect on tissue maceration. Therefore, even higher PelI amounts in the *phoP* mutant can be the reason for its greater maceration ability during the artificial tuber inoculation.

The switch from PehA to PelI as the major pectinolytic enzyme correlates with the most likely sequence of events during plant cell wall polygalacturonate degradation and utilisation by a soft rot pathogen (83). Polygalacturonases have the acidic pH optimum and are therefore required at early stages of infection, as apoplastic fluid is slightly acidic. In contrast, PelI is active at the alkaline pH (81) and is relevant at later stages of infection (when alkalinisation of the infection area occurs).

PelI action would release further amounts of Ca^2+^ and Mg^2+^ and may cause an even stronger decrease of *phoPQ* expression, resulting in the final stabilisation of PhoP regulon members expression at the levels required for the late stages of infection. High Ca^2+^ and Mg^2+^ concentrations would inactivate PhoQ kinase activity, preventing PhoP phosphorylation and blocking PhoP-dependent transcription activation. Contrary to PhoP-dependent transcription activation, PhoP-dependent repression can still occur in these conditions, especially at promoter regions with multiple and/or high affinity PhoP binding sites.

Exhausting homopolygalacturonan amounts (due to its rapid hydrolysis by polygalacturonase) makes high PGA activity unnecessary. Yet, when the infection progresses further, hydrolysis of arabinose containing cell wall components justifies activation of arabinose transporters and enzymes involved in arabinose metabolism.

Many transporters are controlled by PhoP at least to some extent and most of them are activated. High transporter activity at the onset of infection would increase nutrient availability for the pathogen. Increased nutrient availability at the later stages of infection justifies the reduction of transporter gene expression. Citrate transporters seem to be a special case since one of them (CitM) is repressed while the other one (CitW) is activated by PhoP. The switch from CitW to CitM during plant colonisation may correlate with a switch from citrate fermentation to aerobic citrate metabolism. Additional tri- and dicarboxylate transporters can also be controlled by PhoP, but were poorly expressed in the conditions used in our work. Multiple potential PhoP binding sites near other anaerobiosis-related genes justify a dedicated study of PhoP role in anaerobic conditions.

PhoP is also involved in the control of several important stress resistance proteins. Most of their genes are PhoP-activated and hence help *Pve* cells at the early stages of infection. In contrast, a large group of poorly characterised “tellurite resistance” proteins are better expressed at the later stages of infection.

We finally note that the PhoP regulon in *Pve* is likely to be much larger than the 116 DEGs found in this work. We believe that many of the additional PhoP binding sites found by the *in silico* approach are the true ones and can result in PhoP-dependent expression level changes in conditions different from the ones studied here. The list of experimentally established and *in silico* predicted PhoP regulon members determined in the course of this work can be helpful in further studies of regulatory networks and different aspects of *Pve* interaction with its host plants.

## Acknowledgements

This work was supported by grants from Belarusian Republican Foundation for Basic Research and Russian Foundation for Basic Research.

